# Demographic stochasticity can drive population variability even in large populations

**DOI:** 10.64898/2026.10.07.757207

**Authors:** Tetsuya Akita

## Abstract

Demographic stochasticity is generally expected to contribute little to population variability when census abundance is large, because random demographic events are increasingly averaged across many individuals. This expectation, however, depends on how evenly individuals contribute to reproduction. In many natural populations, reproductive success is highly skewed, so that a large census population can coexist with a much smaller effective number of breeders (*N*_b_). Although *N*_b_ is commonly used as a genetic measure of contemporary effective size, it also summarizes variation in reproductive contribution and may therefore contain information about demographic variability. Here, we develop a female-based age-structured theoretical framework linking *N*_b_ to temporal variability in mature population abundance, with *N* denoting the census number of mature females and *N*_b_ the effective number of female breeders. Recruitment variability arises from non-Poisson reproductive variation and environmental stochasticity, and its transmission to mature abundance is described by a life-history coefficient, *γ*, determined by survival, maturation, and the contribution of overlapping cohorts. When the Poisson component of recruitment variance is negligible and *N*_b_*/N* is sufficiently small, the coefficient of variation in mature abundance, CV[*N*], can be approximated from *N*_b_, environmental variability, and *γ*, with little additional information from census abundance. This approximation does not require specification of the functional form of the relationship between adult abundance and recruitment. Stochastic age-structured simulations closely reproduced the analytical prediction, even when additional demographic stochasticity in survival was included. Substantial fluctuations in mature abundance occurred at census sizes on the order of 10^5^ when *N*_b_ was small, whereas overlapping cohorts buffered recruitment fluctuations. Because *N*_b_ can be estimated genetically from samples of a single cohort, the framework provides a direct link between genetic monitoring and population dynamics. More broadly, these results show that large census abundance alone does not guarantee weak demographic variability and highlight the potential of genetic data to inform expected population fluctuations when detailed demographic information is limited.

**Open Research statement:** No new empirical data were generated in this study. The previously published data used in the supplementary analysis are publicly available from Dryad (Clarke et al. 2024; https://doi.org/10.5061/dryad.p2ngf1vzm). Code used to generate the simulation results, analyses, and figures is publicly available at https://github.com/teTUNAakita/DemoStoNbPopVar. If the manuscript is accepted, the version of record of the code will be permanently archived in Zenodo and assigned a DOI.

## 1 Introduction

Temporal fluctuations in population size arise from multiple ecological mechanisms, including environmental variability, density-dependent feedbacks, dispersal and migration, species interactions, and rapid evolutionary responses (e.g., Royama, 1992; Hanski, 1999; Turchin, 2013; Hairston Jr et al., 2005). Demographic stochasticity, reflecting random variation among individuals in survival and reproduction, is another fundamental source of population variability. However, because its relative contribution to population fluctuations is expected to decline as the census number of mature individuals (*N*) increases, demographic stochasticity is often treated as a minor source of variability in large populations (Lande et al., 2003).

This expectation relies on the assumption that reproductive contributions are averaged across many individuals. However, reproductive success can be highly skewed in many organisms. For example, large and old female fish can contribute disproportionately to reproductive output through higher fecundity, longer spawning seasons, or higher offspring quality (Hixon et al., 2014). In highly fecund marine species, sweepstakes reproductive success can further concentrate successful recruitment into a small fraction of adults (Hedgecock and Pudovkin, 2011). In such cases, a large number of mature individuals may be present, while successful recruitment may be generated by only a small fraction of them.

These patterns challenge the use of the census number of mature individuals alone to infer the contribution of demographic stochasticity to population variability. Previous studies have shown that the effects of demographic stochasticity on population dynamics depend strongly on life history in age-structured populations (Jeppsson and Forslund, 2012), and that reproductive heterogeneity can further increase demographic variance (Melbourne and Hastings, 2008). However, how strongly reproductive skew translates into temporal variability in mature abundance remains unclear.

The effective number of breeders (*N*_b_, Waples, 1990) is a measure of contemporary effective size within a single breeding season (e.g., a year), analogous to the effective population size (*N*_e_, Charlesworth, 2009), defined over a generation. Because *N*_b_ decreases as variance in reproductive success among mature individuals, it can be much smaller than the census number of mature individuals when reproductive contributions are highly skewed (Waples, 2024). A recent meta-analysis reported very low *N*_b_*/N* (or *N*_e_*/N*) ratios in many marine species and insects, suggesting that strong reproductive skew may be widespread in taxa with high reproductive output (Clarke et al., 2024). Importantly, *N*_b_ can be estimated from genetic data, for example using sibling or linkage-disequilibrium information from a single cohort, without requiring detailed life-history data (Wang et al., 2016; Caballero et al., 2026). Recently, Akita (2025) developed a theoretical framework linking *N*_b_ to the variation in recruitment and showed that, when both *N*_b_ and *N*_b_*/N* are small, recruitment variability can become substantial and can be approximated from *N*_b_ and environmental variability alone. This raises a further question: given *N*_b_, can temporal variability in the number of mature individuals be predicted from recruitment variability using only limited additional demographic information?

Here, we extend the recruitment-level framework of Akita (2025) to age-structured population dynamics. We derive an analytical relationship between recruitment variability and temporal variability in the number of mature individuals and evaluate it using stochastic age-structured simulations. The analysis separates three major components governing population variability: (i) non-Poisson variation in reproduction and early survival, summarized by *N*_b_; (ii) environmental variability affecting population-level survival; and (iii) a life-history coefficient describing how recruitment fluctuations are transmitted through age structure. The first two components were established previously, whereas the third provides the new link from recruitment variability to variability in mature abundance. We show that when *N*_b_*/N* is sufficiently small, the coefficient of variation (CV[*N*]) can be approximated largely from *N*_b_, environmental variability, and this life-history coefficient, with little additional information from census abundance itself. This result implies that demographic stochasticity associated with reproductive variation can remain an important source of population variability even when census population size is large.

Although the model is described primarily with iteroparous fishes in mind, because these taxa provide particularly suitable applications of the framework, the underlying theory is not restricted to fishes and can be applied more broadly when *N*_b_*/N* is sufficiently small.

## 2 Theory

Previous work established a recruitment model in which recruitment variability is determined by demographic and environmental stochasticity (Akita, 2025). Within this framework, the magnitude of recruitment fluctuations can be expressed as a function of the effective number of breeders and environmental variability (Eqs. 3–4). The first part of the present Theory section briefly summarizes this previously developed framework to establish notation and biological interpretation. We then extend the framework by linking recruitment variability to population variability through age-structured population dynamics. This extension yields an analytical expression for population variability (Eq. 12) and provides a mechanistic link between the effective number of breeders and population variability. The main symbols used in this section are summarized in Table 1.

**Table 1.** List of mathematical symbols used in the main text.

|  |  |
| --- | --- |
| $R$ | Number of female recruits |
| $N$ | Census number of mature females |
| $N_b$ | Effective number of female breeders |
| $Y$ | Total number of female offspring produced after the parental and non-parental processes |
| $P$ | Probability that a female offspring survives to recruitment |
| $\lambda_i$ | Expected reproductive output of mature female $i$ |
| $\phi$ | Overdispersion parameter describing non-parental variation |
| $\sigma_E$ | Log-scale standard deviation of environmentally driven variation in recruitment |
| $\sigma_D$ | Log-scale standard deviation of non-Poisson demographic variation in recruitment |
| $l_a$ | Cumulative survivorship of a female recruit to age $a$ |
| $m_a$ | Maturity at age $a$ |
| $h_a$ | Expected contribution of a female recruit to mature abundance at age $a$ ($= l_a m_a$) |
| $\gamma$ | Life-history coefficient relating recruitment variability to variability in mature abundance |

### 2.1 Conceptual framework

We consider a single closed population with discrete breeding seasons and age-structured population dynamics. Throughout this study, all population quantities are defined for females. In the recruitment-level frameworks of Akita (2020, 2025), the female contribution to recruitment variability was formulated using the effective number of female breeders, *N*_b,f_, whereas *N*_b_ was subsequently used after incorporating the male contribution. Here, because the age-structured model follows females through recruitment, survival, maturation, and reproduction, we use *N*_b_ to denote the effective number of female breeders, corresponding to *N*_b,f_ in the previous formulations. Accordingly, *N* denotes the number of mature females, and offspring and recruits are also restricted to females.

Recruitment variability is a major driver of temporal fluctuations in population size and arises from both demographic and environmental processes. Previous work developed a theoretical framework that partitions this variation into demographic and environmental components and expresses their combined effects as a function of the effective number of breeders and environmental variability. The conceptual structure of this framework is summarized in Fig. 1.

**Figure 1.**
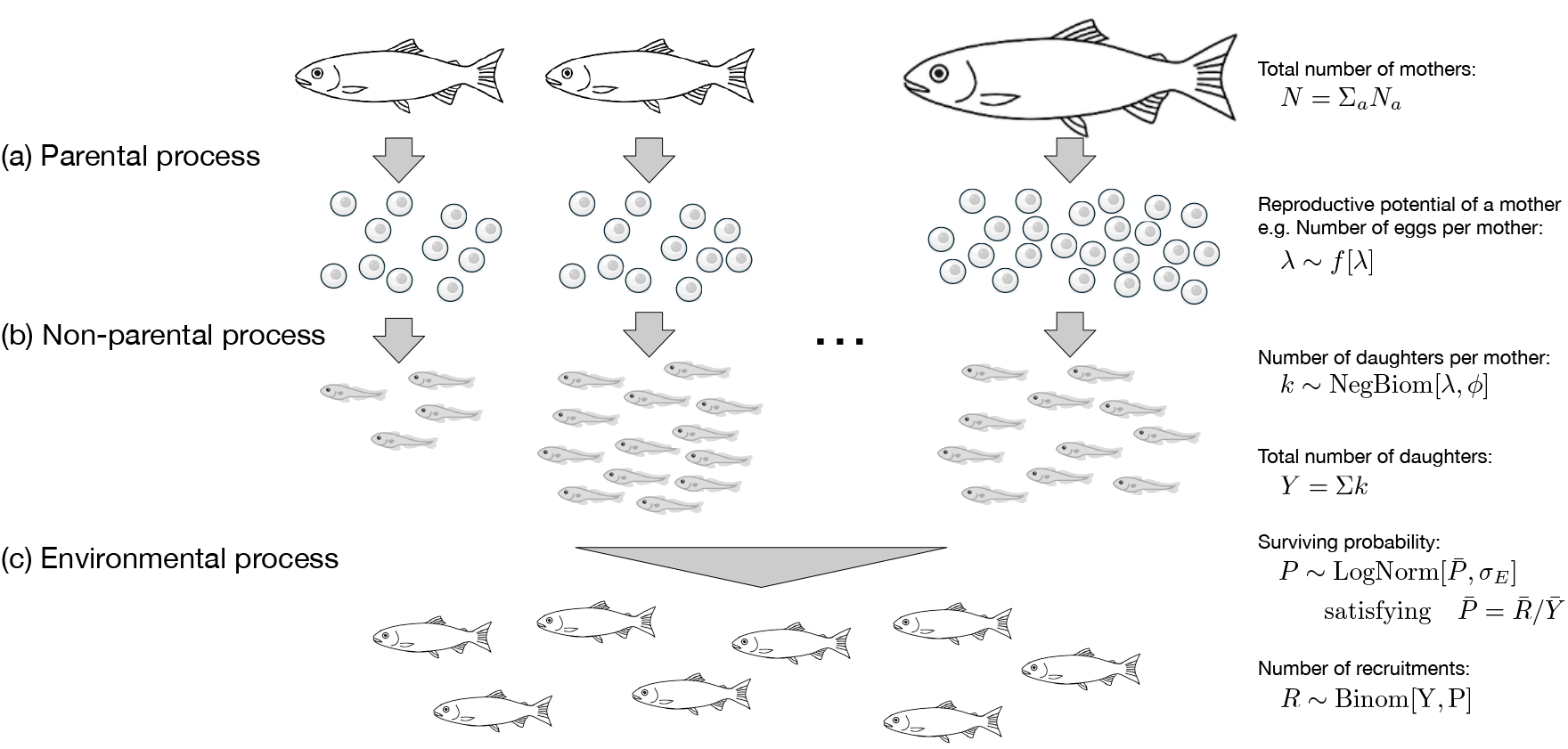
Conceptual framework of the recruitment model. Individual females differ in their expected reproductive output (parental process). Offspring survival exhibits additional family-correlated variation independent of parental reproductive potential (non-parental process). Environmental variability acts at the cohort level after these family-level processes (environmental process). Together, these three processes determine recruitment variability.

The framework consists of three stochastic processes (Fig. 1). Variation among females in expected reproductive output generates the parental component, whereas additional family-correlated variation in female offspring survival generates the non-parental component. Together, these two processes determine the demographic component of recruitment variability, summarized by the effective number of breeders. Environmental processes acting at the cohort level then generate additional variation in recruitment success.

This conceptual framework provides the basis for the recruitment model summarized below, which is then extended to age-structured population dynamics to derive temporal variability in mature abundance.

### 2.2 Recruitment model

We first consider the parental and non-parental processes that together determine the demographic component of recruitment variability (Fig. 1a,b). Parental variation is represented by variation among females in their expected reproductive output, *λ*, summarized by CV[*λ*]. In the non-parental process, offspring survival exhibits additional family-correlated variation that is independent of parental reproductive potential. This process is represented by an overdispersed female offspring distribution with overdispersion parameter *ϕ*, which generates non-parental variation in reproductive success. The combined effects of parental and non-parental variation are summarized by the effective number of female breeders, *N*_b_ (Akita, 2020):

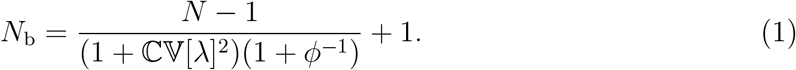

When parental variation is absent (ℂV[*λ*] = 0) and non-parental overdispersion is absent (*ϕ* → ∞), Eq. 1 reduces to *N*_b_ = *N*.

After the parental and non-parental processes have acted, offspring survival is further modified by cohort-level environmental conditions shared among all offspring within a cohort (Fig. 1c). Let 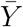 denote the expected total number of female offspring produced after the parental and non-parental processes. Because the expected total reproductive output is 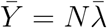, where 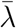 is the mean expected reproductive output among mature females, the expected number of recruits is determined by a density-dependent function, 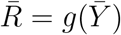, with 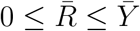. Given 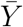 and 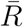, the mean survival probability from female offspring to recruitment is then defined as 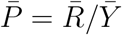.

Environmental stochasticity causes the realized survival probability, *P*, to fluctuate among cohorts around 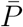. This variation is represented by a lognormal random process with mean 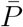 and log-scale standard deviation *σ*_E_. Given the realized number of female offspring, *Y*, the number of female recruits follows

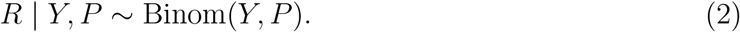

Integrating over variation in *Y* and *P* yields the Poisson–lognormal recruitment process, as shown by Akita (2025).

Under the specified recruitment function, the resulting recruitment variance conditional on *N* is given by

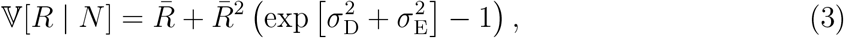

where

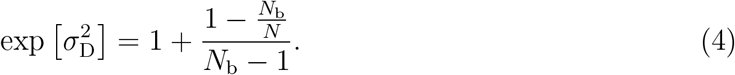

Here, 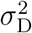 denotes the non-Poisson demographic component of recruitment variance arising from parental and non-parental variation. Equation 4 shows that the magnitude of this component increases as both *N*_b_ and *N*_b_*/N* decrease. The first term, 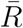, represents Poisson sampling variance in recruitment, whereas the second term represents additional variability arising from non-Poisson demographic stochasticity, environmental stochasticity, and their interaction. Detailed derivations are provided by Akita (2025).

Recruitment variability can be expressed more directly using the coefficient of variation. From Eq. 3,

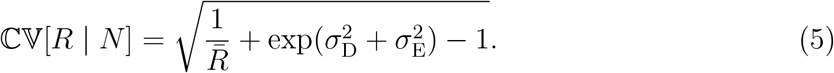

When non-Poisson demographic variability is absent (*σ*_D_ = 0) and environmental stochasticity is negligible (*σ*_E_ = 0), Eq. 5 reduces to the Poisson expectation, 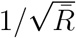. When the Poisson term in Eq. 5 is small relative to the non-Poisson demographic and environmental terms, recruitment variability can be approximated as

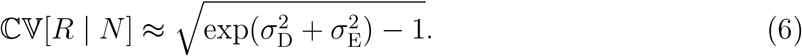

Thus, under the recruitment framework, recruitment variability is governed by the effective number of breeders (*N*_b_), the ratio *N*_b_*/N*, and environmental variability (*σ*_E_). When the Poisson term is negligible, CV[*R* | *N*] is approximately independent of the specific form of the density-dependent function *g*(·) that determines 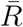.

### 2.3 Linking recruitment variability to population variability

Recruitment variability does not map directly onto population variability because recruitment fluctuations are modified through survival, maturation, and overlapping cohorts. We therefore derive an analytical relationship between recruitment variability and population variability under a discrete-time age-structured population model with annual recruitment. We consider fluctuations around a deterministic equilibrium. Let *R*_*t*_ denote the number of female recruits entering the population in year *t*, and let *N*_*t*_ denote the number of mature females in year *t*. We define *h*_*a*_ = *l*_*a*_*m*_*a*_, where *l*_*a*_ and *m*_*a*_ denote survivorship to age *a* and maturity at age *a*, respectively. Thus, *h*_*a*_ represents the expected contribution of a female recruit to mature abundance at age *a*.

The number of mature females at time *t* can be expressed as the cumulative contribution of recruitment from multiple age classes,

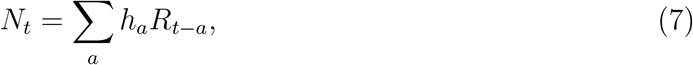

where *R*_*t*−*a*_ denotes recruitment occurring *a* years before year *t*. Equation 7 shows that mature abundance is determined by the accumulated contributions of multiple recruitment cohorts weighted by age-specific survival and maturation schedules.

Assuming that recruitment fluctuations around the deterministic equilibrium are independent among cohorts, the variance in mature female abundance is

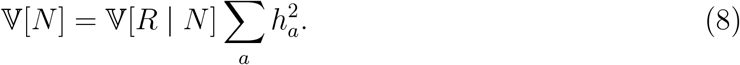

Expressing Eq. 8 in terms of coefficients of variation yields

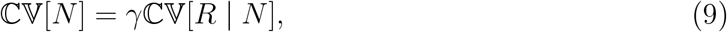

where

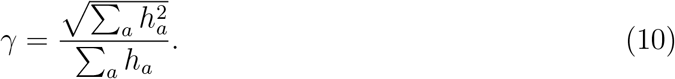

Derivations of Eqs. 8–10 are provided in Appendix S1. The coefficient *γ* summarizes how recruitment fluctuations are transmitted to variability in mature female abundance through age-specific survival and maturation schedules. In this formulation, *γ* is determined by these life-history schedules and is independent of the functional form of recruitment.

If *k* age classes contribute equally to mature abundance (i.e., *h*_*a*_ = *h* for the contributing age classes), Eq. 10 gives 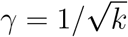, illustrating the reduction in recruitment variability produced by averaging across overlapping cohorts.

A second useful special case is obtained when maturation is complete from a given age onward and annual post-maturation survival is constant at *s*, with no upper limit on age. Because the common survivorship to maturation cancels between the numerator and denominator of Eq. 10, *γ* reduces to

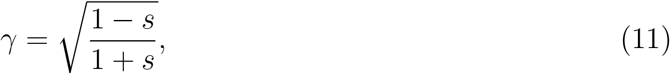

as shown in Appendix S1. Thus, high adult survival reduces the transmission of recruitment fluctuations to mature abundance because multiple overlapping cohorts contribute simultaneously to the population. In contrast, when adult survival is low, mature abundance is concentrated in relatively few recent cohorts, so recruitment fluctuations are transmitted more directly to population variability. This expression has the same mathematical form as that implied by the corresponding special case of Wikström et al. (2012).

Combining Eq. 9 with the expression for recruitment variability derived above yields

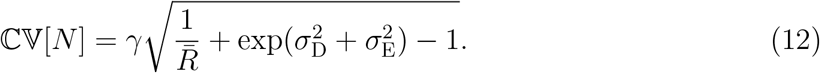

When the Poisson term is small relative to the non-Poisson demographic and environmental components of recruitment variability, population variability can be approximated as

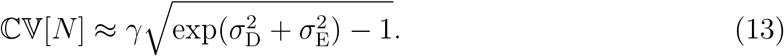

When *N*_b_*/N* is sufficiently small, the dependence of Eq. 4 on *N* becomes negligible. In this regime, ℂV[*R* | *N*] is determined primarily by *N*_b_ and *σ*_E_, and ℂV[*N*] by *N*_b_, *σ*_E_, and *γ*, with little additional information from census abundance itself.

### 2.4 Simulation procedure

We used stochastic age-structured simulations to evaluate the analytical predictions derived above. The simulations followed the same female-only framework as the analytical model, with annual recruitment and age-specific survival and maturation. Unlike the analytical model, in which survival was represented deterministically through the survivorship schedule, survival in the simulations was implemented as a binomial process at each age, thereby allowing additional demographic variation in survival.

Mean recruitment was determined by a Beverton–Holt density-dependent function of total expected reproductive output. In the simulations, individual expected reproductive output, *λ*_*i*_, was represented by body weight, so that 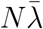 corresponds to spawning stock biomass in the fish-based implementation used here. Mean recruitment was therefore given by

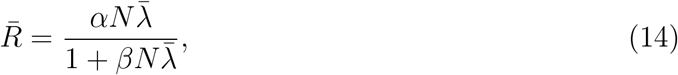

where *α* and *β* determine productivity and the strength of density dependence, respectively. We varied the non-parental overdispersion parameter *ϕ* over a broad range to generate populations with different effective numbers of breeders and *N*_b_*/N* ratios, while keeping the survival, maturation, growth, and stock–recruitment functions fixed. Environmental variability was set to *σ*_E_ = 0, 0.3, or 0.6. Thus, differences among simulation scenarios were generated primarily by variation in reproductive overdispersion and environmental stochasticity.

For each parameter combination, we ran 20 replicate simulations for 10,000 years and discarded the first 500 years as burn-in. We then calculated the mean effective number of breeders, mean mature female abundance, recruitment variability CV[*R*], and mature female population variability CV[*N*]. Simulated values were used to generate representative population trajectories and to compare observed population variability with the analytical predictions from Eqs. 12 and 13. For this life-history schedule, the resulting transmission coefficient was *γ* = 0.58. Additional simulation details and full parameter values are provided in Appendix S2.

## 3 Results

### 3.1 Small *N*_b_ amplifies variability in large populations

Contrasting population trajectories illustrated how reproductive overdispersion and environmental stochasticity affected temporal variability in mature female abundance (Fig. 2). Under a near-Poisson reproductive regime (*ϕ* = 10^2^), the effective number of breeders was close to the census number of mature females (*N*_b_ ≈ 2.2 × 10^5^, *N* ≈ 2.5 × 10^5^, *N*_b_*/N* ≈ 0.9), with the small departure from unity arising from age-dependent differences in reproductive success. When environmental stochasticity was absent (*σ*_E_ = 0), mature female abundance showed only minor temporal fluctuations (Fig. 2a).

**Figure 2.**
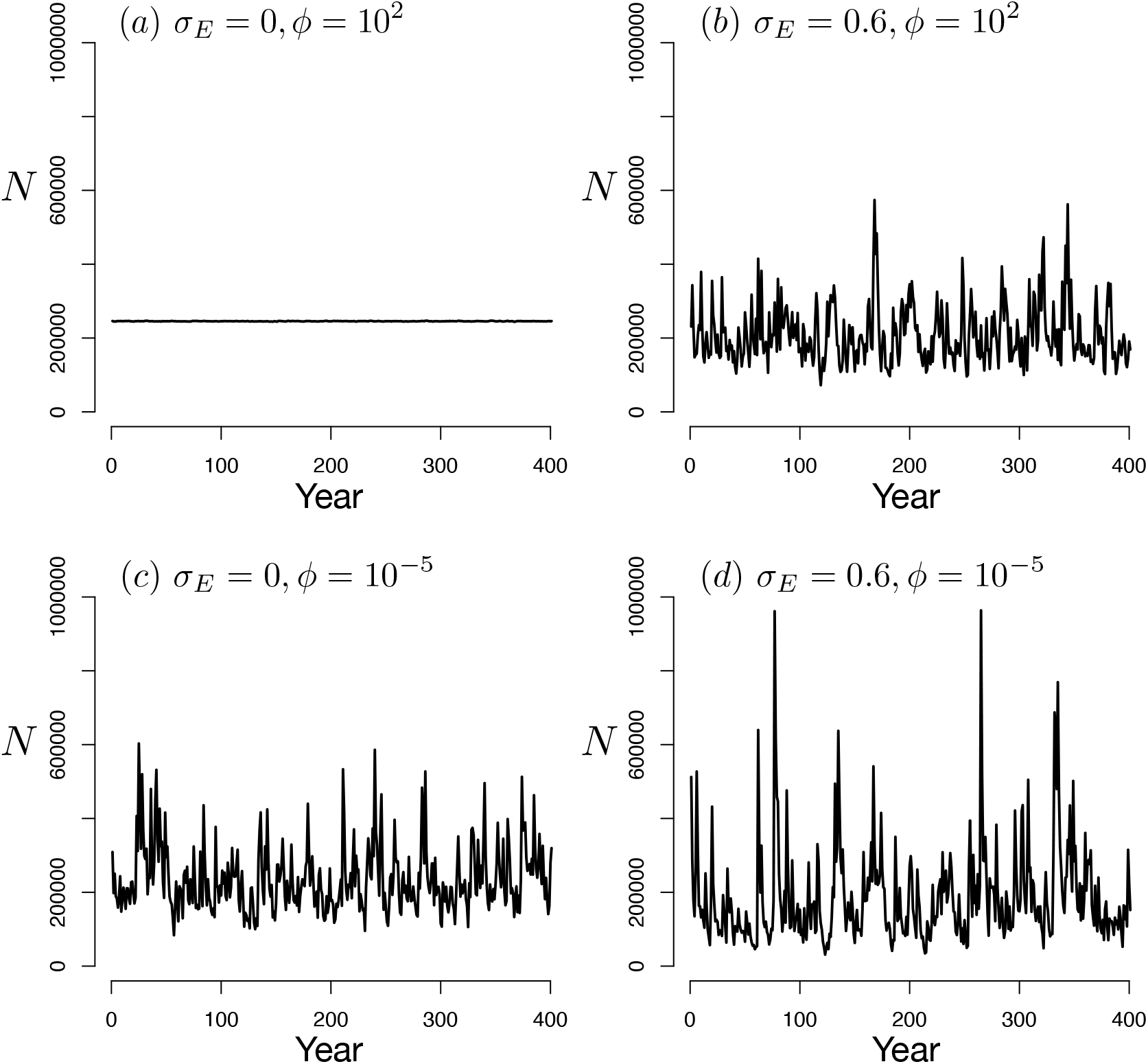
Population trajectories under contrasting reproductive overdispersion and environmental stochasticity. Time series of the census number of mature females (*N*) after burn-in under four parameter combinations: (a) no environmental stochasticity (*σ*_E_ = 0) and weak reproductive overdispersion (*ϕ* = 10^2^); (b) environmental stochasticity (*σ*_E_ = 0.6) and weak reproductive overdispersion; (c) no environmental stochasticity and strong reproductive overdispersion (*ϕ* = 10^−5^); and (d) environmental stochasticity and strong reproductive overdispersion. Each panel shows one illustrative simulation with identical demographic parameters except for *ϕ* and *σ*_E_. Other parameter values are provided in Appendix S2.

Introducing environmental stochasticity (*σ*_E_ = 0.6) under the same reproductive regime substantially increased temporal variability in mature female abundance (Fig. 2b). Thus, when *N*_b_ remained close to *N*, environmental stochasticity was the dominant source of population fluctuations.

By contrast, strong reproductive overdispersion generated pronounced population fluctuations even when environmental stochasticity was absent (Fig. 2c). In this scenario, *N*_b_*/N* declined to approximately 1.5 × 10^−5^, yielding *N*_b_ ≈ 3.5 despite a census population size on the order of 10^5^. Thus, substantial population variability occurred in a numerically large population when the effective number of breeders was very small.

Combining strong reproductive overdispersion with environmental stochasticity produced the largest fluctuations among the four scenarios (Fig. 2d). These trajectories illustrate that a large census population does not necessarily imply low population variability when reproductive skew reduces *N*_b_ to very small values, and that environmental stochasticity can further amplify these fluctuations.

### 3.2 Population variability increases as the effective number of breeders declines

We next quantified how temporal variability in mature female abundance changed with *N*_b_ across simulations in which mature female abundance remained on the order of 10^5^ (Fig. 3). For each simulation, we calculated the coefficient of variation in mature female abundance, CV[*N*], after burn-in and compared the simulated values with the analytical approximation in Eq. 13.

**Figure 3.**
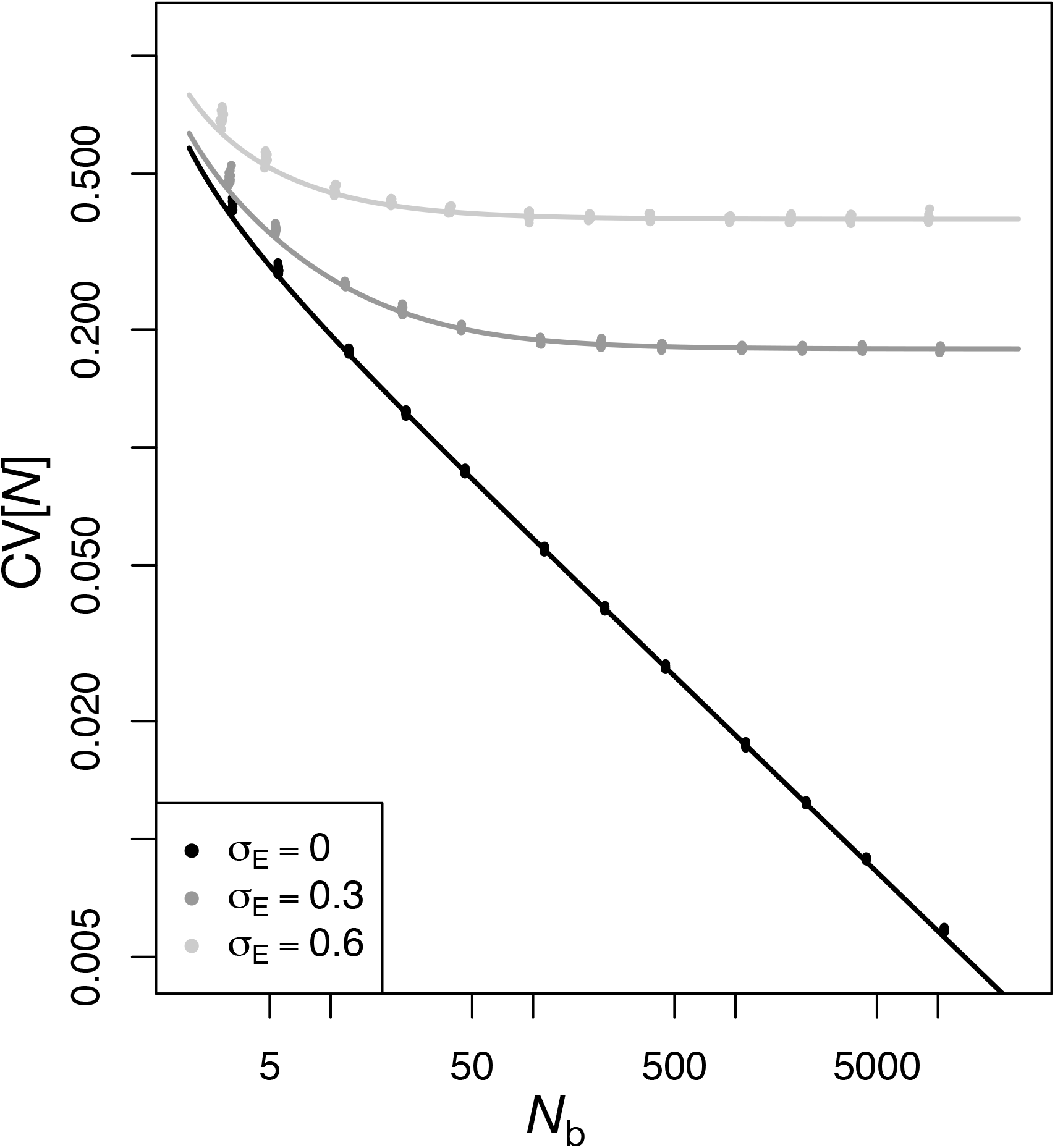
Relationship between the effective number of breeders, *N*_b_, and temporal variability in mature female abundance, CV[*N*], in the stochastic age-structured simulations. Points show values calculated from individual simulation replicates after the burn-in period. Lines show the analytical approximation in Eq. 13, which neglects the Poisson term and assumes that *N*_b_*/N* is sufficiently small. Results are shown for *σ*_E_ = 0, 0.3, and 0.6. The life-history schedule used in the simulations corresponds to *γ* = 0.58.

Population variability increased as the effective number of breeders declined. In the absence of environmental stochasticity (*σ*_E_ = 0), CV[*N*] remained low at large *N*_b_ but increased sharply as *N*_b_ became small. Thus, strong reproductive overdispersion generated substantial temporal variability in mature female abundance even when census abundance remained large.

Environmental stochasticity increased population variability across the range of *N*_b_. At large *N*_b_, CV[*N*] approached a level determined primarily by environmental variability, whereas at smaller *N*_b_, population variability increased progressively above this level as the non-Poisson demographic component became more important.

The analytical approximation closely reproduced the simulated relationship between *N*_b_ and CV[*N*] across the environmental conditions considered (Fig. 3). The approximation remained accurate despite the additional demographic variation in survival included in the simulations, which was not represented explicitly in the analytical model. This agreement indicates that, when *N*_b_*/N* is sufficiently small, population variability can be approximated from *N*_b_, *σ*_E_, and the life-history coefficient *γ*, with little additional information from census abundance. The corresponding relationship between *N*_b_ and recruitment variability is shown in Fig. S2.

## 4 Discussion

The present study extends the demographic interpretation of the effective number of breeders developed by Akita (2025) from recruitment variability to temporal variability in mature population abundance. Whereas the previous framework showed that *N*_b_ can summarize the magnitude of non-Poisson recruitment variability when *N*_b_*/N* is sufficiently small, the present analysis identifies the additional demographic information required to translate this variability to an age-structured population. Specifically, the life-history coefficient *γ* summarizes how survival, maturation, and overlapping cohorts modify recruitment fluctuations, allowing population variability to be approximated from *N*_b_, environmental variability, and relatively limited life-history information.

### 4.1 Reproductive skew can decouple census abundance from demographic variability

A common expectation in population ecology is that the relative contribution of demographic stochasticity declines as population size increases, because random outcomes of many approximately independent demographic events are averaged over larger numbers of individuals (e.g., Lande et al., 2003; Engen et al., 2010; Jeppsson and Forslund, 2012). This conventional sampling component typically declines in proportion to 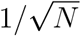 when demographic outcomes arise from approximately independent contributions of *N* individuals. In our framework, the corresponding Poisson sampling component scales with 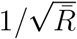. Our results show, however, that the same intuition does not necessarily apply to variability generated by reproductive skew. When *N*_b_*/N* is sufficiently small, the non-Poisson component becomes only weakly dependent on census abundance and mean recruitment, as reflected in the approximation in Eq. 13.

The contrast between survival and reproduction is informative in this regard. In the simulations, survival was implemented as a binomial process and therefore introduced an additional source of demographic stochasticity that was not represented explicitly in the analytical model. Nevertheless, this contribution was small at the large census abundances considered here, and the analytical approximation remained accurate. By contrast, recruitment-driven variability remained substantial when the effective number of breeders, *N*_b_, was small. Thus, a population can be large enough for conventional sampling variation in survival to be negligible while still experiencing pronounced demographic fluctuations generated by reproductive skew.

This difference arises because census abundance and effective breeder number summarize different aspects of the population. Census abundance determines the magnitude of individual-level sampling variation in demographic processes such as survival, whereas *N*_b_ summarizes the extent to which reproductive success is concentrated among mature individuals. When *N*_b_*/N* is sufficiently small, the non-Poisson component of recruitment variability becomes only weakly dependent on *N* and is governed primarily by *N*_b_ (Akita, 2025). Consequently, a large number of mature individuals does not necessarily imply weak demographic variability when highly uneven reproductive success results in a small effective number of breeders.

### 4.2 Predicting population variability from limited demographic information

A central implication of the present framework is that, under strong reproductive skew, temporal variability in mature abundance can be predicted from a small number of aggregate quantities. Population variability in age-structured species may in principle depend on many demographic features, including reproductive heterogeneity, survival, maturation, density dependence, and environmental variation. When *N*_b_*/N* is sufficiently small, however, much of this complexity is summarized by *N*_b_, which captures non-Poisson variation in reproductive contribution; *σ*_E_, which describes cohort-level environmental variation in recruitment; and *γ*, which describes how recruitment fluctuations are transmitted through survival and maturation.

Existing approaches to stochastic demography derive demographic variance from age- or stage-specific survival and reproductive rates, together with information on their variability, obtained either from repeated individual-level observations or from distributional assumptions (e.g., Engen et al., 2005, 2009; Caswell and Vindenes, 2018). In contrast, *N*_b_ summarizes reproductive heterogeneity at the population level and can be estimated genetically without separately observing the demographic processes that generated that heterogeneity. This reduction may be particularly useful in data-limited populations, where detailed estimates of multiple vital rates and their temporal variances are unavailable.

The use of effective-size estimates in population monitoring has expanded in recent years, including their incorporation into broader biodiversity-monitoring frameworks (e.g., Hoban et al., 2020; Mergeay et al., 2026). These applications have primarily emphasized the genetic consequences of small effective size, such as genetic drift, loss of genetic variation, and inbreeding. The present framework highlights a complementary demographic interpretation of *N*_b_. Because *N*_b_ reflects variation in reproductive contribution within a breeding season (Waples, 2024), it can therefore contain information about stochastic variation generated during reproduction (Akita, 2025). This dual interpretation is particularly relevant because *N*_b_ can be estimated genetically from single-cohort samples (Bertram et al., 2024; Robinson et al., 2024). Genetic monitoring may therefore provide information not only about genetic status but also about expected demographic variability.

Combined with approximate information on environmental variability and basic survival and maturation schedules, genetically estimated *N*_b_ may allow expected population variability to be constrained before a complete age- or stage-structured demographic model can be constructed. Importantly, the approximate expression for CV[*N*] (Eq. 13) does not require specification of the functional form or local slope of the relationship between adult abundance and recruitment. Instead, recruitment variability is summarized by *N*_b_ and *σ*_E_, while its transmission to mature abundance is summarized by *γ*. In some cases, plausible ranges for *σ*_E_ and *γ* could be informed by better-studied populations or species with similar environments and life histories, providing an initial estimate of expected population variability when detailed demographic information is unavailable.

### 4.3 How age structure buffers recruitment variability

Age structure can substantially alter how fluctuations generated during recruitment appear in mature population abundance. When several cohorts coexist in the mature population, cohort-specific recruitment anomalies are averaged across ages, whereas populations dominated by a small number of cohorts retain more of the temporal signal generated at recruitment. This buffering effect is captured by *γ* in the present framework.

Previous studies have shown that age structure can modify the population-level consequences of stochastic recruitment in several ways. Bjørnstad et al. (2004) showed that recruitment fluctuations can be distributed across multiple adult cohorts, generating cohort resonance and low-frequency variability in aggregate stock abundance. Worden et al. (2010) further showed that this response depends on life history, with reduced adult survival strengthening cohort resonance. Similarly, Legendre et al. (1999) found that demographic stochasticity associated with reproduction had stronger effects in short-lived than in long-lived birds, because individuals surviving across years buffered fluctuations in the number of descendants. Wikström et al. (2012) demonstrated that age truncation and endogenous recruitment feedback can further modify the transmission of recruitment variability to adult abundance. Like the age-structured analyses of Bjørnstad et al. (2004); Worden et al. (2010); Wikström et al. (2012), our analysis considers fluctuations around a deterministic equilibrium. However, rather than explicitly modeling local feedback from adult abundance to subsequent recruitment, the present framework treats recruitment deviations around equilibrium as stochastic inputs and focuses on their transmission through survivorship and maturation schedules.

An important ecological implication of this separation is that similar recruitment variability need not translate into similar variability in mature abundance. In the present framework, *N*_b_ and *σ*_E_ characterize the magnitude of variability generated during recruitment, whereas *γ* determines how strongly that variability is expressed in mature abundance through survival and maturation schedules. Consequently, populations experiencing similar recruitment fluctuations may show markedly different population variability because of differences in their life histories.

### 4.4 Empirical relevance of low *N*_b_*/N* **and small** *N*_b_

The extremely small values of the non-parental overdispersion parameter *ϕ* used in some simulations were intended to illustrate the limiting behavior of the model rather than to represent typical parameter values in natural populations. The empirical relevance of the framework should therefore not be evaluated from *ϕ* itself, because a given value of *N*_b_*/N* can arise from different combinations of non-parental overdispersion and variation among mature individuals in expected reproductive output. More directly relevant quantities are the effective-to-census ratio, which determines when the dependence on census abundance becomes weak, and the absolute value of *N*_b_, which determines the magnitude of the non-Poisson component of demographic variability.

Low effective-to-census size ratios have been reported particularly frequently in highly fecund taxa characterized by high early mortality, including many marine organisms and invertebrates (Clarke et al., 2024). Such patterns are consistent with the long-standing concept of sweepstakes reproductive success, in which only a small fraction of potential breeders contribute disproportionately to successful recruitment (Hedgecock and Pudovkin, 2011). However, a low *N*_b_*/N* ratio alone does not imply strong demographic variability: if absolute *N*_b_ remains large, the non-Poisson contribution generated by reproductive heterogeneity may still be small.

To examine whether low effective-to-census ratios tend to coincide with small absolute *N*_b_, we re-summarized the non-replicated dataset of Clarke et al. (2024) using populations for which both *N*_b_ and census size (denoted by *N*_c_) were available (Appendix S3). Among populations with *N*_b_*/N*_c_ *<* 0.1, 72.4% also had *N*_b_ *<* 100. Because the empirical *N*_b_ estimates generally combine both sexes and *N*_c_ does not necessarily correspond to the number of mature females represented by *N* in our model, this comparison should be regarded as an empirical analogue rather than a direct quantitative validation of the theoretical relationship.

These two quantities play different roles in the theory. A sufficiently small *N*_b_*/N* ratio defines the regime in which the dependence of recruitment variability on census abundance becomes weak, whereas the absolute magnitude of *N*_b_ determines the strength of the non-Poisson component of recruitment variability. Previous analysis suggested that *N*_b_*/N <* 0.1 provides a useful practical regime for the approximation (Akita, 2025), but this threshold should not be interpreted as a boundary below which demographic variability necessarily becomes large. Rather, substantial recruitment-driven variability is expected when a low ratio is accompanied by a sufficiently small absolute *N*_b_. The re-analysis of Clarke et al. (2024) indicates that these two conditions can frequently coincide in empirical populations. Because taxonomic representation in the available dataset was highly uneven (Table S2 in Appendix S3), broader comparative datasets will be required to determine how generally low *N*_b_*/N* and small absolute *N*_b_ coincide across taxa.

### 4.5 Limitations and future extensions

The present framework treats age-specific survivorship deterministically, whereas environmental variation may also affect juvenile and adult survival. Incorporating stochastic survival would introduce additional sources of variability after recruitment, potentially together with temporal autocorrelation and covariance among age-specific vital rates. In such cases, population variability would generally no longer be summarized by a single transmission coefficient such as *γ*.

The present formulation also defines *N*_b_ as the effective number of female breeders, whereas empirical genetic estimates of *N*_b_ generally incorporate reproductive contributions from both sexes. Extending the framework to two-sex populations would therefore require explicit treatment of sex ratio, mating structure, and sex-specific variation in reproductive success. These extensions would increase biological realism, but would not alter the central conclusion that large census abundance alone does not guarantee low population variability when reproductive contribution is highly uneven and *N*_b_ remains small.

## Acknowledgement

This work was supported by JSPS KAKENHI Grant Number 23K05945, 25K02038, and 26K01761.

