## Supplemental materials for "Demographic stochasticity can drive population variability even in large populations"

#### Appendix S1

**Authors:** Tetsuya Akita

**Manuscript title:** *Demographic stochasticity can drive population variability even in large populations*

### Appendix S1. Derivation of the relationship between recruitment variability and population variability

We derive the relationship between temporal variability in recruitment and temporal variability in mature female abundance under the age-structured population model described in the main text. Let $R_t$ denote the number of female recruits entering the population in year $t$, and let $N_t$ denote the number of mature females in year $t$. We consider fluctuations around a deterministic equilibrium and assume that recruitment fluctuations are independent among cohorts. Thus,

$$\mathbb{E}[R_t] = \bar{R}, \tag{A1}$$

$$\mathbb{V}[R_t \mid N] = \mathbb{V}[R \mid N], \tag{A2}$$

for all $t$.

Let

$$h_a = l_a m_a, \tag{A3}$$

where $l_a$ is cumulative survivorship to age $a$ and $m_a$ is maturity at age $a$. The quantity $h_a$ therefore represents the expected contribution of a female recruit to mature abundance at age $a$. Mature female abundance at time $t$ is

$$N_t = \sum_a h_a R_{t-a}. \tag{A4}$$

Taking expectations of Eq. (A4),

$$\mathbb{E}[N_t] = \mathbb{E}\left[\sum_a h_a R_{t-a}\right] \tag{A5}$$

$$= \sum_a h_a \mathbb{E}[R_{t-a}] \tag{A6}$$

$$= \bar{R} \sum_a h_a. \tag{A7}$$

Hence, the equilibrium mean abundance of mature females is

$$\bar{N} = \bar{R} \sum_a h_a. \tag{A8}$$

The variance of mature female abundance is

$$\mathbb{V}[N_t] = \mathbb{V}\left[\sum_a h_a R_{t-a}\right] \tag{A9}$$

$$= \sum_a h_a^2 \mathbb{V}[R_{t-a} \mid N] + 2 \sum_{a<b} h_a h_b \operatorname{Cov}(R_{t-a}, R_{t-b} \mid N). \tag{A10}$$

Under the assumption that recruitment fluctuations around the deterministic equilibrium are independent among cohorts,

$$\mathrm{Cov}(R_{t-a}, R_{t-b} \mid N) = 0 \qquad (a \neq b), \tag{A11}$$

and therefore

$$\mathbb{V}[N_t] = \sum_a h_a^2 \mathbb{V}[R_{t-a} \mid N] \tag{A12}$$

$$= \mathbb{V}[R \mid N] \sum_a h_a^2. \tag{A13}$$

Using Eqs. (A8) and (A13), the coefficient of variation in mature female abundance is

$$\mathbb{CV}[N] = \frac{\sqrt{\mathbb{V}[N]}}{\bar{N}} \tag{A14}$$

$$= \frac{\sqrt{\mathbb{V}[R \mid N] \sum_a h_a^2}}{\bar{R} \sum_a h_a} \tag{A15}$$

$$= \frac{\sqrt{\sum_a h_a^2}}{\sum_a h_a} \frac{\sqrt{\mathbb{V}[R \mid N]}}{\bar{R}}. \tag{A16}$$

Because

$$\mathbb{CV}[R \mid N] = \frac{\sqrt{\mathbb{V}[R \mid N]}}{\bar{R}}, \tag{A17}$$

we obtain

$$\mathbb{CV}[N] = \gamma \, \mathbb{CV}[R \mid N], \tag{A18}$$

where

$$\gamma = \frac{\sqrt{\sum_a h_a^2}}{\sum_a h_a}. \tag{A19}$$

Thus, $\gamma$ summarizes the extent to which recruitment variability is transmitted to variability in mature female abundance through age-specific survivorship and maturation schedules.

A simple closed-form expression for $\gamma$ can be obtained when maturation is complete from age $a_m$ onward and post-maturation annual survival is constant at $s$. In this case,

$$h_{a_m+k} = l_{a_m} s^k, \qquad k = 0, 1, 2, \ldots \tag{A20}$$

so that

$$\sum_{k=0}^{\infty} h_{a_m+k} = \frac{l_{a_m}}{1-s}, \tag{A21}$$

$$\sum_{k=0}^{\infty} h_{a_m+k}^2 = \frac{l_{a_m}^2}{1-s^2}. \tag{A22}$$

Substituting these expressions into the definition of $\gamma$ gives

$$\gamma = \frac{l_{a_m}/\sqrt{1-s^2}}{l_{a_m}/(1-s)} = \sqrt{\frac{1-s}{1+s}}. \tag{A23}$$

The common survivorship to maturation, $l_{a_m}$, cancels, so under this special case the transmission of recruitment variability depends only on post-maturation survival. As $s$ increases, recruitment fluctuations are increasingly averaged across overlapping cohorts, whereas at low $s$ mature abundance is dominated by fewer recent cohorts and recruitment fluctuations are transmitted more directly.

This special case is closely related to previous analytical results for stochastic age-structured populations. Wikström et al. (2012) derived the stationary variability of adult abundance in a stage-structured model with stochastic recruitment and endogenous recruitment feedback, resulting in an AR(2) process. When recruitment feedback is absent, their expression implies a ratio of adult to recruitment coefficients of variation of $\sqrt{(1-s)/(1+s)}$ under constant adult survival, identical to the special case of $\gamma$ derived above.

#### Appendix S2. Simulation details and parameter values

We used stochastic age-structured simulations to evaluate the analytical relationship between recruitment variability and temporal variability in mature female abundance. The simulations followed the same female-only framework as the analytical model. Recruitment occurred annually, and the population was divided into 10 age classes (ages 0–9). Females matured at age 3, such that maturity was 0 for ages 0–2 and 1 for ages 3–9. Annual survival probability was fixed at 0.5 for all age classes.

Individual expected reproductive output, $\lambda_i$, was represented by age-specific body weight, and $N$ denotes the census number of mature females. Relative body weight at age $a$ was derived from the von Bertalanffy growth function for length, combined with the allometric relationship between body weight and length:

$$w_a = \{1 - \exp[-K(a - a_0)]\}^b, \tag{B1}$$

with $K = 0.25$, $a_0 = -1.5$, and $b = 3$. Because expected reproductive output was represented by body weight, total expected reproductive output, $N\bar{\lambda}$, corresponded to spawning stock biomass in this fish-based implementation.

Mean recruitment was determined by a Beverton–Holt density-dependent recruitment function,

$$\bar{R} = \frac{\alpha N\bar{\lambda}}{1 + \beta N\bar{\lambda}}, \tag{B2}$$

where $\alpha = 2000$ and $\beta = 0.001$.

Spawning biomass per recruit (SPR) was calculated from the age-specific survivorship, maturity, and body-weight schedules as

$$\mathrm{SPR} = \sum_a l_a m_a w_a, \tag{B3}$$

where $l_a$ and $m_a$ denote survivorship and maturity at age $a$, respectively. Because $N\bar{\lambda}$ corresponds to spawning stock biomass, the replacement line is given by

$$\bar{R} = \frac{N\bar{\lambda}}{\mathrm{SPR}}. \tag{B4}$$

To illustrate the deterministic equilibrium of the simulation model, we plotted the Beverton–Holt recruitment function together with the replacement line (Fig. S1). Their intersection defines the deterministic equilibrium of the population.

In the analytical model, age-specific survival was treated deterministically through the survivorship schedule, $l_a$. In the simulations, survival was instead implemented as a binomial process to evaluate whether additional demographic variation arising from survival affected the analytical approximation. Because this source of variation scales inversely with population size, its contribution was expected to be small at the population sizes considered here.

For a given year, the effective number of female breeders was calculated from the realized age composition and age-specific expected reproductive output. Let $f_a$ denote the proportion of mature females in age class $a$. Then,

$$\bar{\lambda} = \sum_a f_a w_a, \tag{B5}$$

$$\overline{\lambda^2} = \sum_a f_a w_a^2, \tag{B6}$$

and

$$N_{\mathrm{b}} = \frac{N-1}{(1+\phi^{-1})\overline{\lambda^2}/\bar{\lambda}^2} + 1. \tag{B7}$$

The non-parental overdispersion parameter $\phi$ was varied over

$10^{-5}$, $2\times10^{-5}$, $5\times10^{-5}$, $10^{-4}$, $2\times10^{-4}$, $5\times10^{-4}$, $10^{-3}$, $2\times10^{-3}$, $5\times10^{-3}$, $10^{-2}$, $2\times10^{-2}$, $5\times10^{-2}$.

Environmental variability was set to $\sigma_{\mathrm{E}} = 0$, 0.3, or 0.6.

Conditional on $N$, $N_{\mathrm{b}}$, and $\bar{R}$, recruitment variance was

$$\mathbb{V}[R \mid N] = \bar{R} + \bar{R}^2 \left[\exp(\sigma_{\mathrm{E}}^2)\left(1 + \frac{1 - N_{\mathrm{b}}/N}{N_{\mathrm{b}} - 1}\right) - 1\right]. \tag{B8}$$

Realized recruitment was generated from a Poisson–lognormal distribution parameterized to have mean $\bar{R}$ and the variance given above.

Each simulation was run for 10,000 annual time steps. The initial population consisted of 10,000 age-0 females and no individuals in older age classes. The first 500 years were discarded as burn-in. Twenty replicate simulations were conducted for each combination of $\phi$ and $\sigma_{\mathrm{E}}$. A simulation was treated as extinct only when no individuals remained in any age class.

For each non-extinct replicate, we calculated the mean effective number of female breeders, mean mature female abundance, and the sample coefficients of variation in recruitment and mature female abundance over the post-burn-in period.

For comparison with the simulations, we used the analytical approximation obtained by neglecting the Poisson term and assuming that $N_{\mathrm{b}}/N$ was sufficiently small. Recruitment variability was therefore approximated as

$$\mathbb{CV}[R] \approx \sqrt{\exp(\sigma_{\mathrm{E}}^2)\left(1 + \frac{1}{N_{\mathrm{b}} - 1}\right) - 1}, \tag{B9}$$

and mature female population variability as

$$\mathbb{CV}[N] \approx \gamma\, \mathbb{CV}[R], \tag{B10}$$

where

$$\gamma = \frac{\sqrt{\sum_a h_a^2}}{\sum_a h_a}, \qquad h_a = l_a m_a. \tag{B11}$$

The corresponding relationships between $N_{\mathrm{b}}$ and $\mathbb{CV}[N]$ and between $N_{\mathrm{b}}$ and $\mathbb{CV}[R]$ are shown in Fig. 2 and Fig. S2, respectively.

### Appendix S3. Empirical co-occurrence of low $N_{\mathrm{b}}/N$ and small $N_{\mathrm{b}}$

To examine whether low effective-to-census ratios tend to coincide with small absolute effective breeder numbers in empirical populations, we re-summarized the publicly available dataset compiled by Clarke et al. (2024). We used the non-replicated dataset, in which spatial and temporal replication had been removed, so that repeatedly sampled populations did not receive disproportionate weight in the summary.

We retained records for which both an estimate of the effective number of breeders ($N_\mathrm{b}$) and census size ($N_\mathrm{c}$) were available. Records with missing census size or with census size reported only as a lower bound (e.g., $> 5000$ or $> 6500$) were excluded because an exact $N_\mathrm{b}/N_\mathrm{c}$ ratio could not be calculated. We then calculated the ratio directly as

$$N_\mathrm{b}/N_\mathrm{c} = \frac{N_\mathrm{b}}{N_\mathrm{c}}, \tag{B12}$$

rather than relying on the ratio field provided in the database. This procedure yielded 77 records with both $N_\mathrm{b}$ and $N_\mathrm{c}$ available.

The frequency of $N_\mathrm{b}/N_\mathrm{c} < 0.1$ differed among the taxonomic groups represented in the dataset (Table S2). The criterion was met by 19 of 37 freshwater-fish populations, four of 12 amphibian populations, all three invertebrate populations, two of four mammal populations, and one of two marine-fish populations, whereas none of the 19 diadromous-fish populations met the criterion. These taxon-specific proportions should be interpreted cautiously because sample sizes were highly unequal and were particularly small for several groups.

We summarized the absolute magnitude of $N_\mathrm{b}$ among populations with low $N_\mathrm{b}/N_\mathrm{c}$ ratios. Among the 29 populations with $N_\mathrm{b}/N_\mathrm{c} < 0.1$, 13 (44.8%) had $N_\mathrm{b} < 50$, 21 (72.4%) had $N_\mathrm{b} < 100$, and 24 (82.8%) had $N_\mathrm{b} < 200$. When a more restrictive ratio threshold of $N_\mathrm{b}/N_\mathrm{c} < 0.05$ was used, 7 of 13 populations (53.8%) had $N_\mathrm{b} < 50$, 11 (84.6%) had $N_\mathrm{b} < 100$, and 12 (92.3%) had $N_\mathrm{b} < 200$. For $N_\mathrm{b}/N_\mathrm{c} < 0.02$, the corresponding numbers were 3 of 4 populations (75.0%) for each of the three $N_\mathrm{b}$ thresholds.

These summaries were intended only to assess whether a low effective-to-census ratio can empirically coincide with a small absolute effective breeder number. The thresholds of $N_\mathrm{b} = 50$, 100, and 200 were therefore used as descriptive reference values rather than as biological thresholds. In particular, the present theory does not predict a universal critical value of $N_\mathrm{b}$ at which demographic variability becomes large.

Two differences between the empirical compilation and our theoretical model should also be noted. First, although the database contains many estimates of generational effective population size ($N_\mathrm{e}$), we did not include these because the present framework is formulated specifically in terms of the effective number of breeders contributing to a breeding season. Second, the $N_\mathrm{b}$ estimates in the comparative dataset are study-specific genetic estimates and were not standardized to the effective number of female breeders, whereas $N_\mathrm{b}$ in our theoretical model is defined for female breeders only. Because $N_\mathrm{c}$ in the comparative dataset does not necessarily correspond to the number of mature females represented by $N$ in our model, $N_\mathrm{b}/N_\mathrm{c}$ should be regarded as an empirical analogue rather than an exact counterpart of the theoretical $N_\mathrm{b}/N$ ratio.

Table S1: Parameter values used in the stochastic age-structured simulations.

| Parameter | Description | Value(s) |
|---|---|---|
| Number of age classes | Ages included in the model | 10 (ages 0–9) |
| Age at maturity | First mature age | 3 |
| Annual survival | Survival probability at each age | 0.5 |
| $K$ | Growth coefficient | 0.25 |
| $a_0$ | Growth-curve parameter | $-1.5$ |
| $b$ | Weight-scaling exponent | 3 |
| $\alpha$ | Beverton–Holt productivity parameter | 2000 |
| $\beta$ | Beverton–Holt density-dependence parameter | 0.001 |
| $\sigma_{\mathrm{E}}$ | Environmental variability | 0, 0.3, 0.6 |
| $\phi$ | Non-parental overdispersion | $10^{-5} \sim 5 \times 10^{-2}$ (see text) |
| Simulation length | Number of annual time steps | 10,000 |
| Burn-in | Initial years discarded | 500 |
| Replicates | Simulations per parameter combination | 20 |
| Initial abundance | Number of age-0 females | 10,000 |

Table S2: Frequency of populations with $N_{\mathrm{b}}/N_{\mathrm{c}} < 0.1$ among taxonomic groups in the non-replicated dataset.

| Taxonomic group | Total $n$ | $N_{\mathrm{b}}/N_{\mathrm{c}} < 0.1$ | Percent |
|---|---|---|---|
| Invertebrate | 3 | 3 | 100.0 |
| Freshwater fish | 37 | 19 | 51.4 |
| Mammal | 4 | 2 | 50.0 |
| Marine fish | 2 | 1 | 50.0 |
| Amphibian | 12 | 4 | 33.3 |
| Diadromous fish | 19 | 0 | 0.0 |

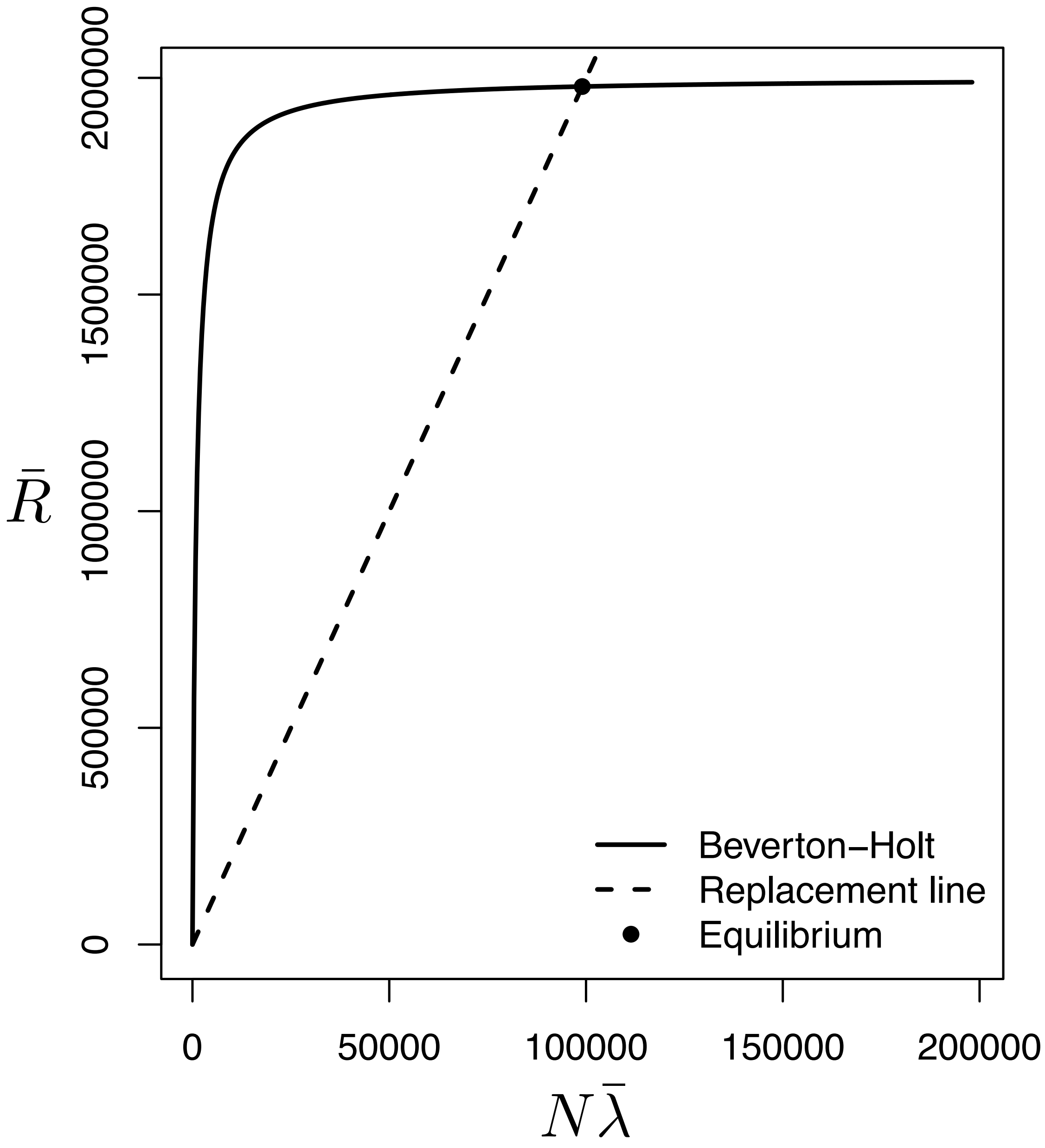


Figure S1: Beverton–Holt recruitment function and replacement line used to define the deterministic equilibrium of the simulation model. The solid curve shows expected recruitment as a function of total expected reproductive output, $N\bar{\lambda}$, and the dashed line shows the replacement line determined by spawning biomass per recruit (SPR). Their intersection, indicated by the point, represents the deterministic equilibrium.

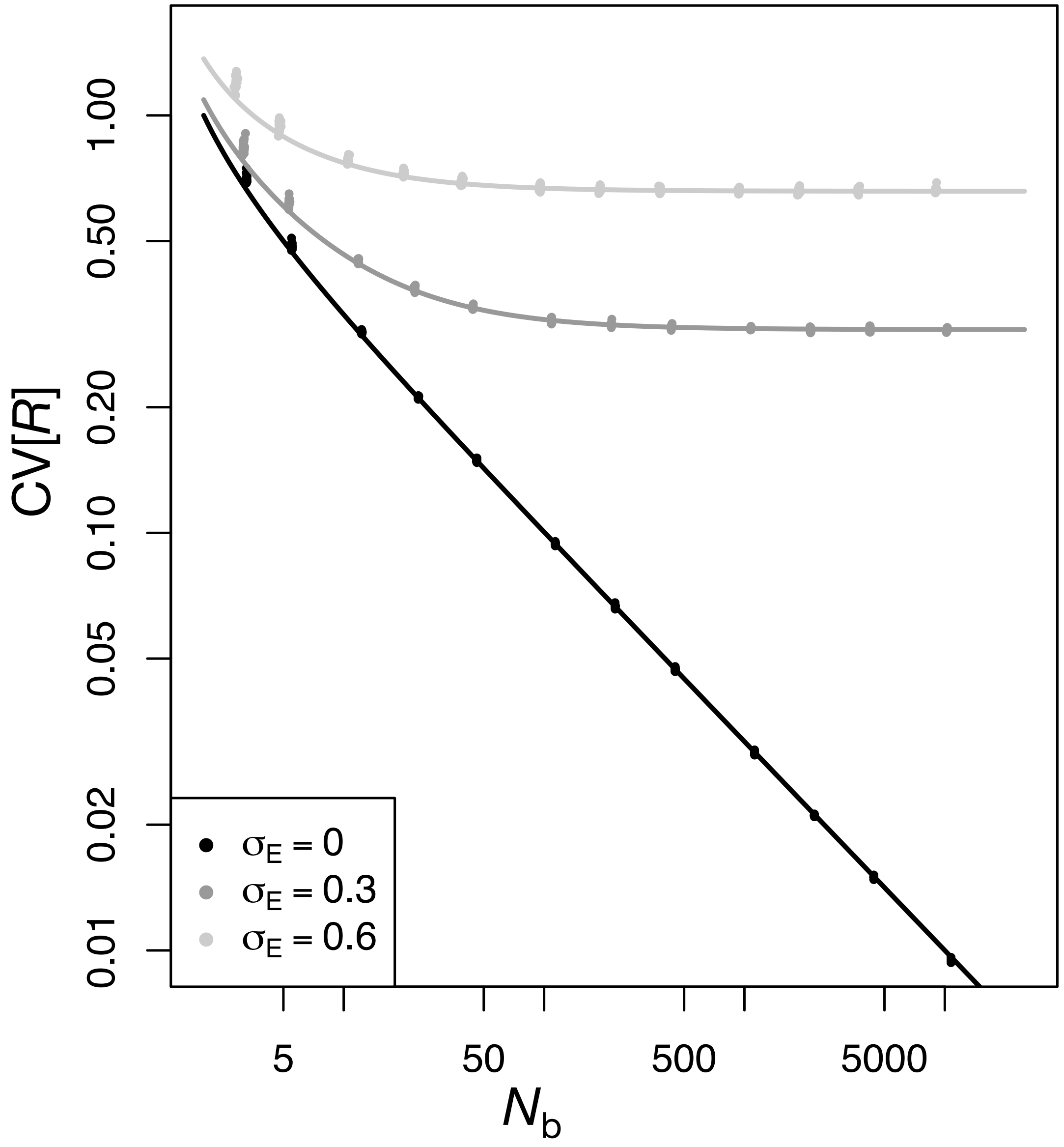


Figure S2: Relationship between the effective number of female breeders, $N_{\mathrm{b}}$, and recruitment variability, $\mathbb{CV}[R]$, obtained from the stochastic age-structured simulations. Points show values calculated from individual simulation replicates after the burn-in period. Lines show the analytical approximation obtained by neglecting the Poisson term and the dependence on $N_{\mathrm{b}}/N$. Results are shown for $\sigma_{\mathrm{E}} = 0$, 0.3, and 0.6.
